## Supplementary Information for "Expanding the toolbox: Novel class IIb microcins show activity against Gram-negative ESKAPE and plant pathogens"

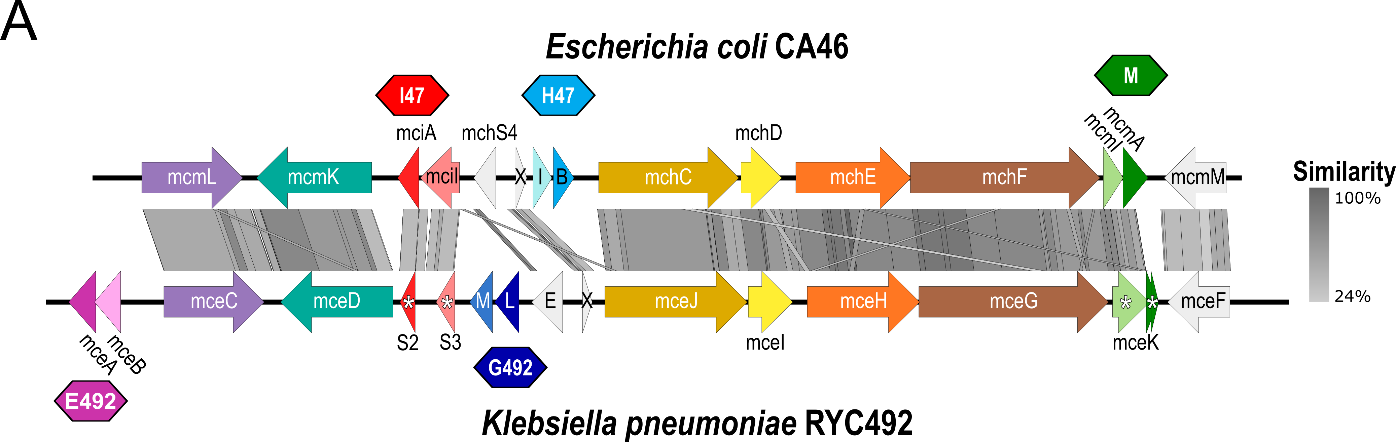

**Fig S1: Comparison of class IIb microcin gene clusters from Escherichia coli CA46 and Klebsiella pneumoniae RYC492.** BLAST sequence comparison of gene clusters was created using Easyfig[1]. Antimicrobial and immunity genes of each cluster are represented by darker and lighter shades, respectively. Colored polygons represent functional microcins MccE492 (pink) and MccG492 (dark blue) from K. pneumoniae as well as MccH47 (light blue), MccI47 (red), and MccM (green) from E. coli. X=mchX, I=mchI, B=mchB, E=mceE, L=mceL, M=mceM. Asterisks indicate truncated and non-functional microcin or immunity genes.

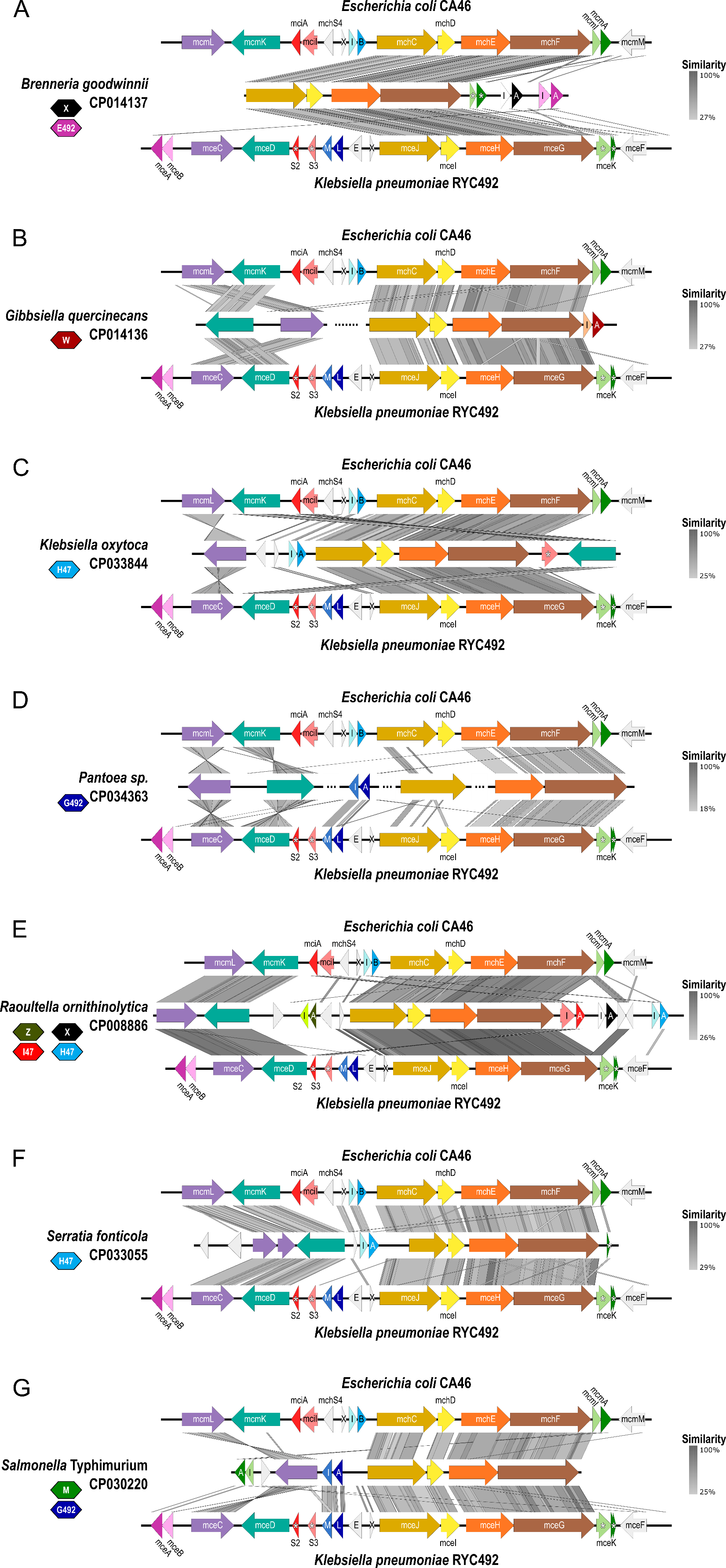

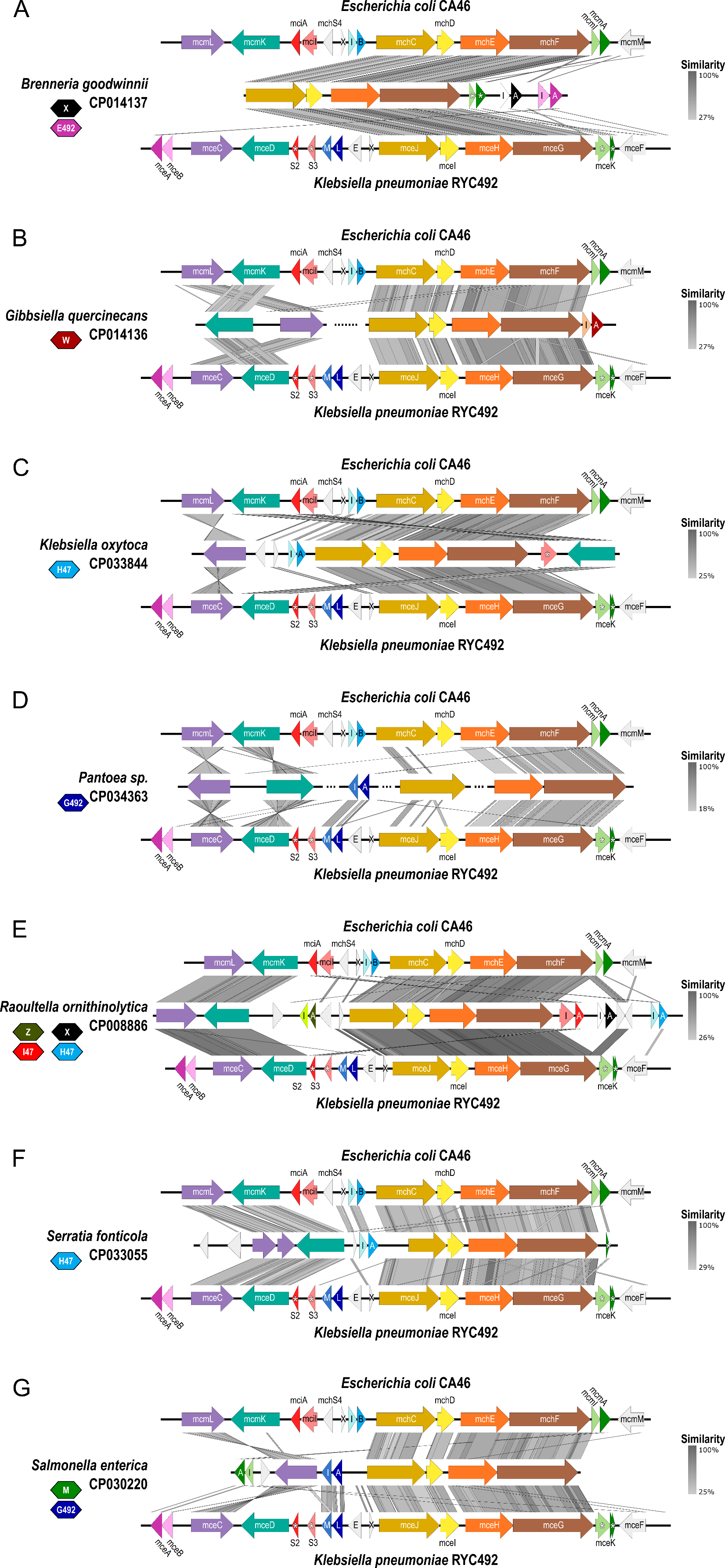

**Fig S2: Comparison of class IIb microcin gene cluster from newly identified genomes with Escherichia coli CA46 and Klebsiella pneumoniae RYC492.** BLAST sequence comparison of gene clusters was created using Easyfig[1]. Antimicrobial and immunity genes of each cluster are represented by darker and lighter shades, respectively. Colored polygons represent novel microcins from the genome indicated on the left: MccE492 (pink), MccG492 (dark blue), MccH47 (light blue), MccI47 (red), MccM (green), MccX (black), MccW (dark red), and MccZ (dark green). X=mchX, I=mchI, B=mchB, E=mceE, L=mceL, M=mceM. Asterisks indicate truncated and non-functional microcin or immunity genes.

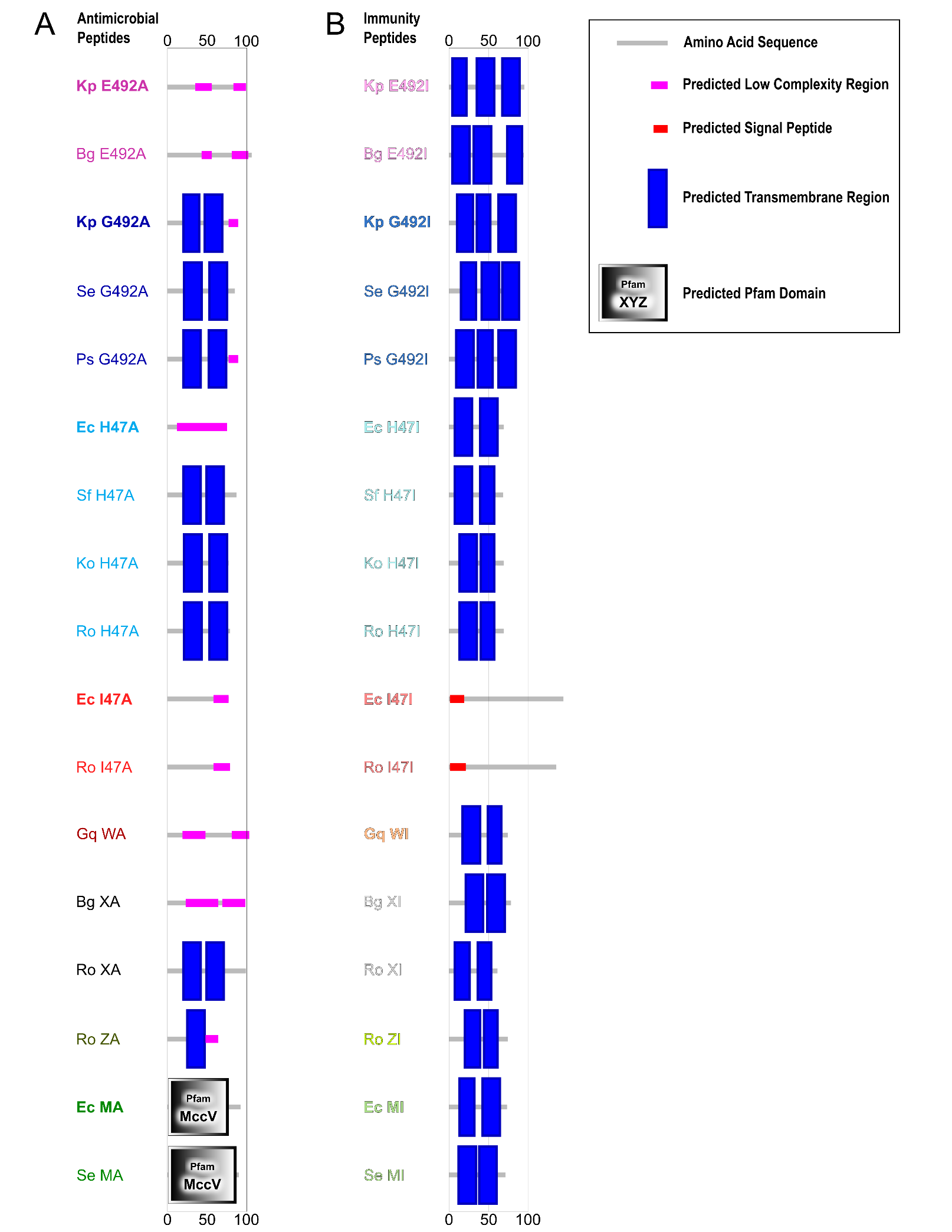

**Fig S3: Region and domain prediction of antimicrobial and immunity peptides using SMART**[2]**.** Previously known proteins are highlighted in bold.
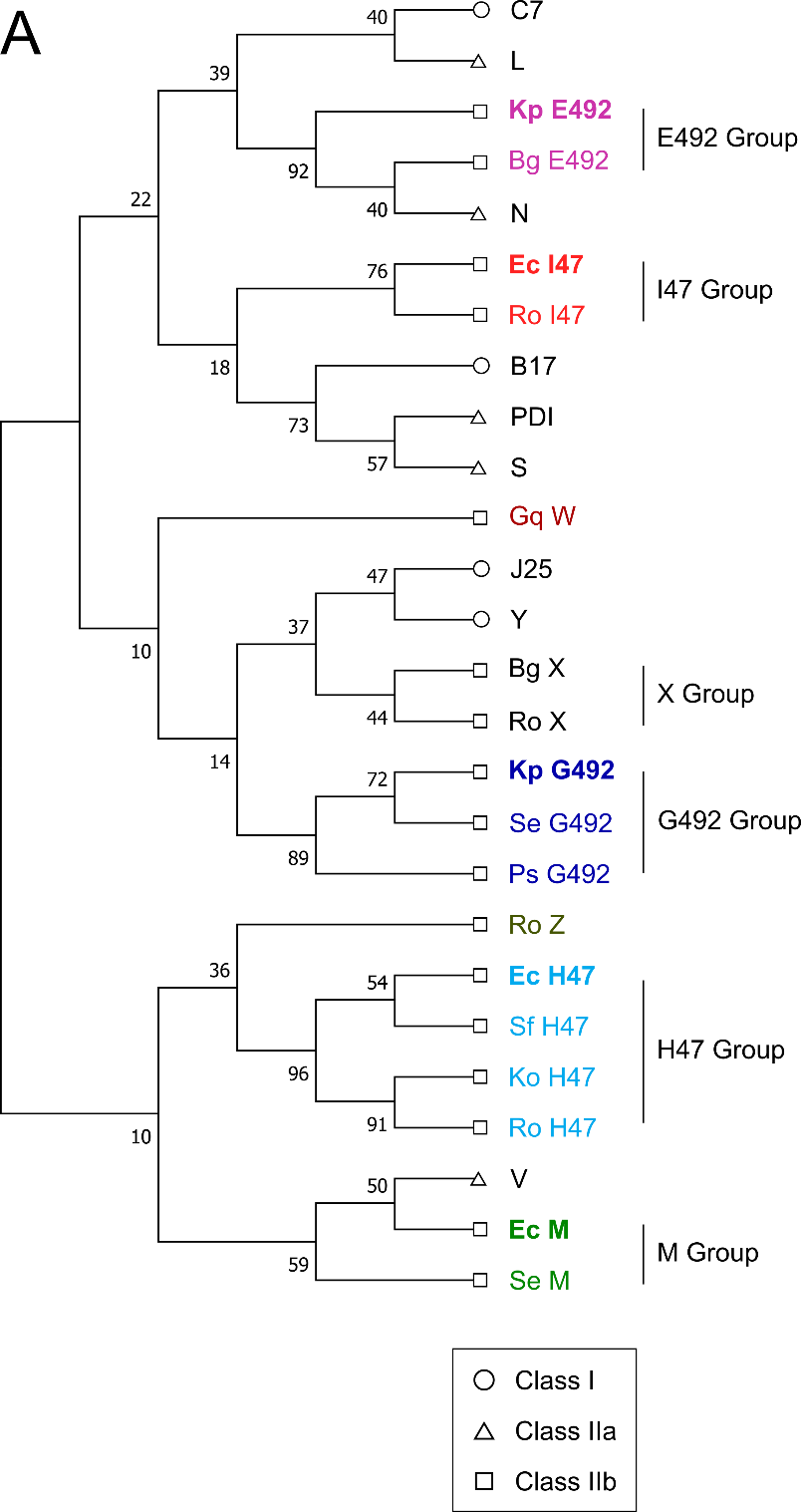

**Fig S4: Phylogeny of all microcins from the classes I, IIa, and IIb.** Phylogenetic tree of all known microcin genes using codon-aligned nucleotide sequences with General Time Reversible model with discrete gamma distribution (GTR+G). The respective signal peptides were removed to ensure alignment of the active regions with antimicrobial activity. Previously known class IIb microcins are highlighted in bold.

**Table S1: Blastp results and closest matches to the known or novel class IIb microcins.** Red color indicates no significant match found.

| **Microcin name** | **Species** | **Accession no.** | **Antimicrobial Gene Name** | **Antimicrobial closest global match** | **Identical/Total Length** | **E-value** | **Immunity Gene Name** | **Immunity closest global match** | **Identical/Total Length** | **E-value** |
| --- | --- | --- | --- | --- | --- | --- | --- | --- | --- | --- |
| *Bg* X | *Brenneria goodwinii* | CP014137 | *Bg* XA | *Ro* XA | 55/98 | 2.0E-12 | *Bg* XI | *Ro* XI | 29/78 | 4.0E-14 |
| *Gq* W | *Gibbsiella quercinecans* | CP014136 | *Gq* WA | *Bg* E492A | 24/96 | 3.7E-01 | *Gq* WI | *Se* MI | 24/74 | 2.5E-01 |
| *Ro* Z | *Raoultella ornithinolytica* | CP008886 | *Ro* ZA | *Ec* MA (mcmA) | 21/64 | 1.0E-03 | *Ro* ZI | *Sf* H47I | 12/74 | 4.1E-01 |
| *Ro* X | *Raoultella ornithinolytica* | CP008886 | *Ro* XA | *Bg* XA | 52/99 | 4.0E-13 | *Ro* XI | *Bg* XI | 29/61 | 4.0E-07 |

**Supplementary Information References**

1.	Sullivan MJ, Petty NK, Beatson SA. Easyfig: a genome comparison visualizer. *Bioinformatics* 2011; **27**: 1009–1010.

2.	Letunic I, Khedkar S, Bork P. SMART: Recent updates, new developments and status in 2020. *Nucleic Acids Res* 2021; **49**: D458–D460.
